# Complete Transcriptome Profiling of Normal and Age-Related Macular Degeneration Eye Tissues Reveals Changes in Regulation of Non-Coding RNA and Extreme Disregulation of Anti-Sense Transcription

**DOI:** 10.1101/143917

**Authors:** Eun Ji Kim, Gregory G. Grant, Anita S. Bowman, Naqi Haider, Harini V. Gudiseva, Venkata Ramana Murthy Chavali

## Abstract

Strand specific RNA sequencing of retina and RPE-Choroid-Scleara (RCS) in age-related macular degeneration (AMD) and matched normal controls reveals striking impact on anti-sense transcription and changes in the regulation of non-coding RNA that has not previously been reported. Hundreds of genes, which do not express anti-sense transcripts in normal retina and RCS, demonstrate extreme anti-sense expression in AMD. And conversely anti-sense transcription is completely abrogated in many genes which express a high level of anti-sense transcripts in normal retina and RCS. Several pathways are very highly enriched in the upregulated anti-sense transcripts - in particular the EIF2 signaling pathway. These results call for a deeper investigation into anti-sense and noncoding RNA regulation in AMD and their potential as therapeutic targets.

## Introduction

Age-related macular degeneration (AMD) is the third largest cause of vision loss worldwide^1^. It is a progressive retinal disorder that involves loss of central vision, hypo-and hyper-pigmentation of the RPE, deposition of drusen in the Bruch’s membrane, and loss of photoreceptors, especially in the 8^th^ or 9^th^ decade^2-4^. The most severe visual loss due to AMD occurs when the disease progresses to one of the two advanced forms: dry (atrophic) AMD or wet (exudative or neovascular) AMD. Genome-Wide Association studies (GWAS) have implicated the role of mutations/variations in the genes involved in various biological pathways such as complement and immune system, cholesterol metabolism, collagen/extra-cellular matrix processing and angiogenesis with onset, progression, and involvement of different stages of AMD ^5-11^. However, most of these reported variations occur in the protein coding regions of genes which account for only 1.5% of the entire human genome^12^. With the development of next-generation sequencing technologies, it has become increasingly apparent that most of the non-protein coding regions of the genome encode for non-coding RNAs (ncRNAs). The ncRNA in eukaryotes are produced in large numbers and exceed the total number of protein-coding genes^13^.

Transcriptome studies have generated significant interest in the role of ncRNAs in the maintenance of cellular processes and function. Based on their length, ncRNAs are broadly divided into short ncRNAs (<200 nucleotides: comprising ribosomal RNA (rRNA)), small interfering RNA (siRNA), micro RNA (miRNA), small nuclear RNA (snRNA), small nucleolar RNA (snoRNA), piwi interacting RNA (piRNA) and long ncRNA (lncRNA, >200 nucleotides)^14^. They may be antisense, intergenic, interleaved, or overlapping with protein-coding genes ^15-17^. In particular, their ability to base pair with other transcripts suggests they may be responsible for a variety of regulatory functions^18^. Transcriptome studies over the last two decades analyzed the posterior region of the eye using SAGE, microarrays and RNA Sequencing methodologies^19-21^. However, none of these studies have analyzed/compared the differences in the transcriptome expression between the normal and AMD retinal tissues.

The posterior part of the eye consists of neural retina, RPE and choroid layers. The RPE secretes a variety of growth factors to help maintain the structural integrity of choriocapillaris and photoreceptors. It also phagocytoses the photoreceptors and regulates ion and metabolic transport between the retina and choroid ^22^. The macula is the cone rich, central part of the retina that is responsible for central vision and is affected in AMD. The retinal photoreceptors, RPE, and choroid act synergistically to maintain visual function, making these tissues logical targets for transcriptome studies of AMD. Yoshida *et al*.^12^ first reported Gene expression studies on young and elderly human retinas, using a slide microarray. Elevated expression levels for *KIAA0120, TRPIP1,* and *ISGF3G* genes were observed in the younger retina when compared to the elderly retina ^23^. Several microarray transcriptome studies on young vs. old, and fetal vs. adult total retina or macular retina identified enriched genes in the fovea macula and peripheral retina ^24, 25^, highlighting the effect of topological location on gene expression ^26^. The differentially expressed genes in macular retina, peripheral retina and enriched RPE were analyzed on a SAGE platform^27^. These studies lead to the identification of tissue and regional specificity of retinal gene expression and assessment of alternative transcription in these tissues^27^.

Normal human retina and RCS were also sequenced using the RNA-Seq platform to measure absolute gene expression levels, identify unknown transcripts, and quantify transcripts of coding and noncoding RNA that may not be defined through custom microarray hybridization and serial analysis of gene expression (SAGE) techniques^21, 28, 29^. Unlike the retinal samples, transcriptome analysis of the RPE-Choroid and RPE-Choroid-Scleara (RCS) tissue layers were pursued together due to difficulty in separating individual layers in cadaver donor eye tissues without contamination. Access to high quality tissues allowed for a comprehensive transcriptome profiling of the retina and RCS. Many unannotated non-coding genes exist in introns as anti-sense to the parent gene; therefore they can be more easily identified by strand specific sequencing. Surprisingly, however, strand-specific sequencing revealed a startling amount of differential anti-sense transcription of protein coding genes with significant enrichment of several pathways. This extreme level of differential expression suggests potential functional and clinical relevance of anti-sense transcription in AMD. In accordance with these findings, the focus of this paper is two-fold: non-coding RNA and anti-sense transcription.

A rapidly growing body of evidence points to the importance of anti-sense and noncoding RNAs in normal and a wide variety of pathological processes. Our study is the first to report a comprehensive transcriptome analysis of the total non-coding and anti-sense RNA profiles in the Retina and RCS tissues of normal and AMD donor eyes. Our results indicate that anti-sense and noncoding RNAs may play an important role in the development and progression of AMD.

## Results

### Analysis of gene expression

Hierarchical clustering based on Jensen-Shannon divergence was performed for the sense and antisense gene expression, as well as only the noncoding RNA (Fig.1). We observed a clear delineation between the PR and PRCS tissues considering sense expression, for both coding and noncoding regions. The separation within AMD and normal tissue types, however, is not apparent. Using the antisense gene expression, we observed a clear clustering of the samples by their tissue type and disease status. We have analyzed all three transcript types (gene-sense, gene-anti-sense and noncoding) in our study.

**Fig.1.**
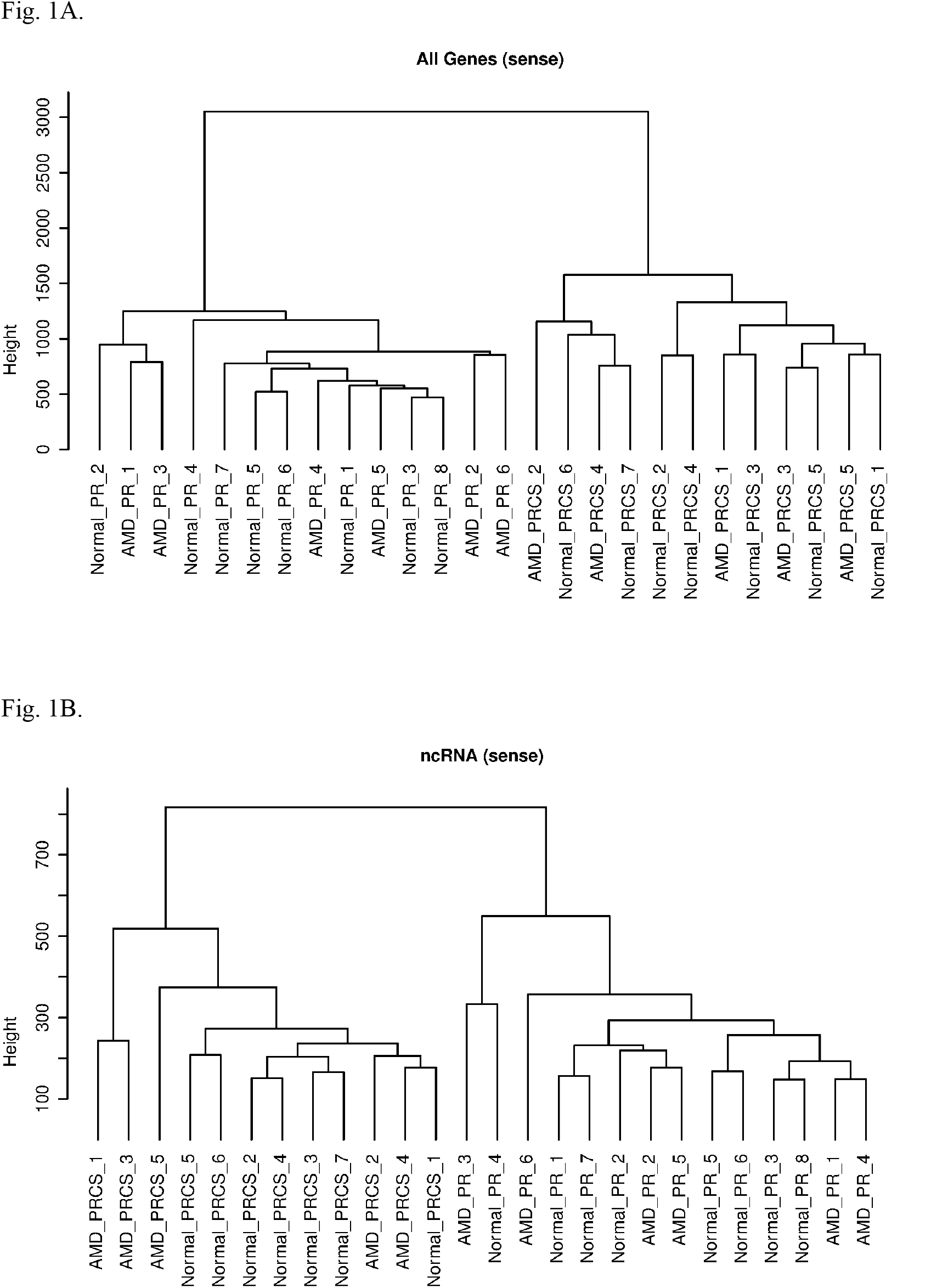
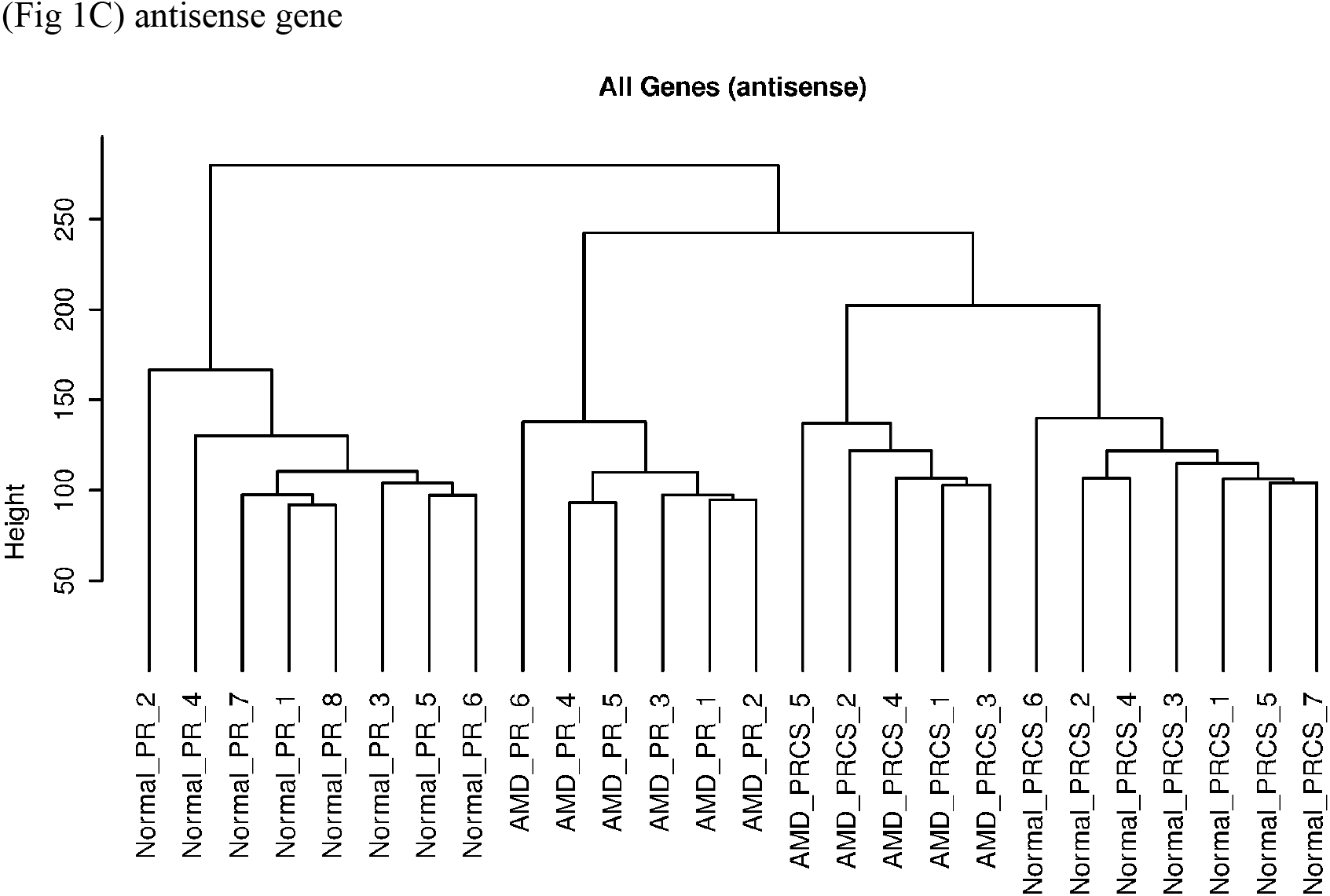
Hierarchical clustering of genes across 26 samples. (A) Clustering of sense gene expression, coding and noncoding. (B) Clustering of only noncoding genes. (C) Clustering of antisense gene expression

Expression of ncRNAs, defined as a gene having PORT normalized average coverage ≥ 1 was determined. In total, 6,972 ncRNA were expressed in at least one sample using this criterion. The average number of expressed ncRNA is 3,582 in AMD-PR, 3210 in AMD-PRCS, 3725 in normal PR, and 3,330 in normal PRCS. We quantified the number of various biotypes of noncoding among those expressed (Fig.2B). The vast majority of expressed ncRNA comprised of antisense and lincRNA. The absence of the smaller noncoding types is due in part to the size selection in the library construction process. In both of the aforementioned categories however, the number of expressed transcripts is highest in Normal-PR.

**Fig. 2.**
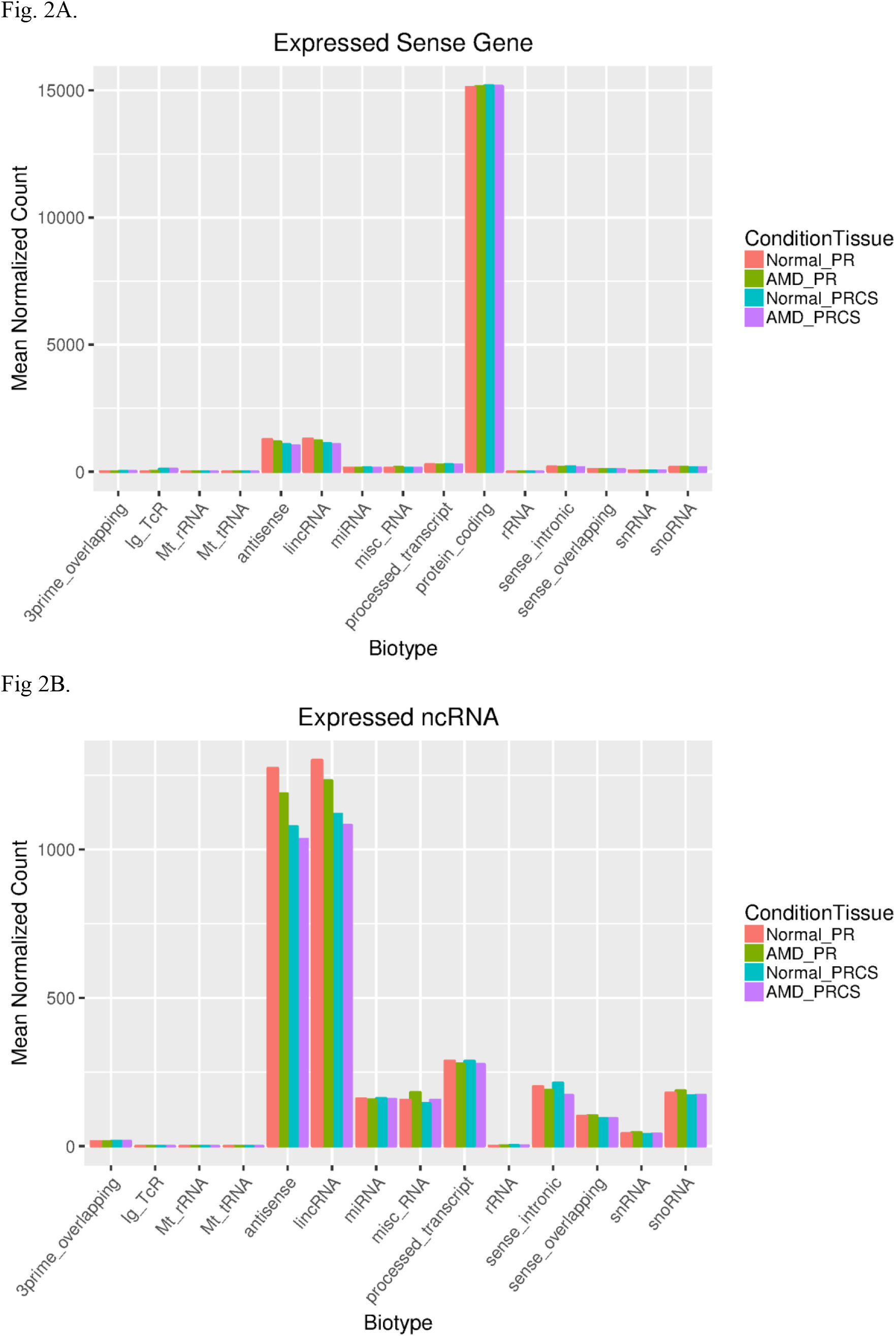
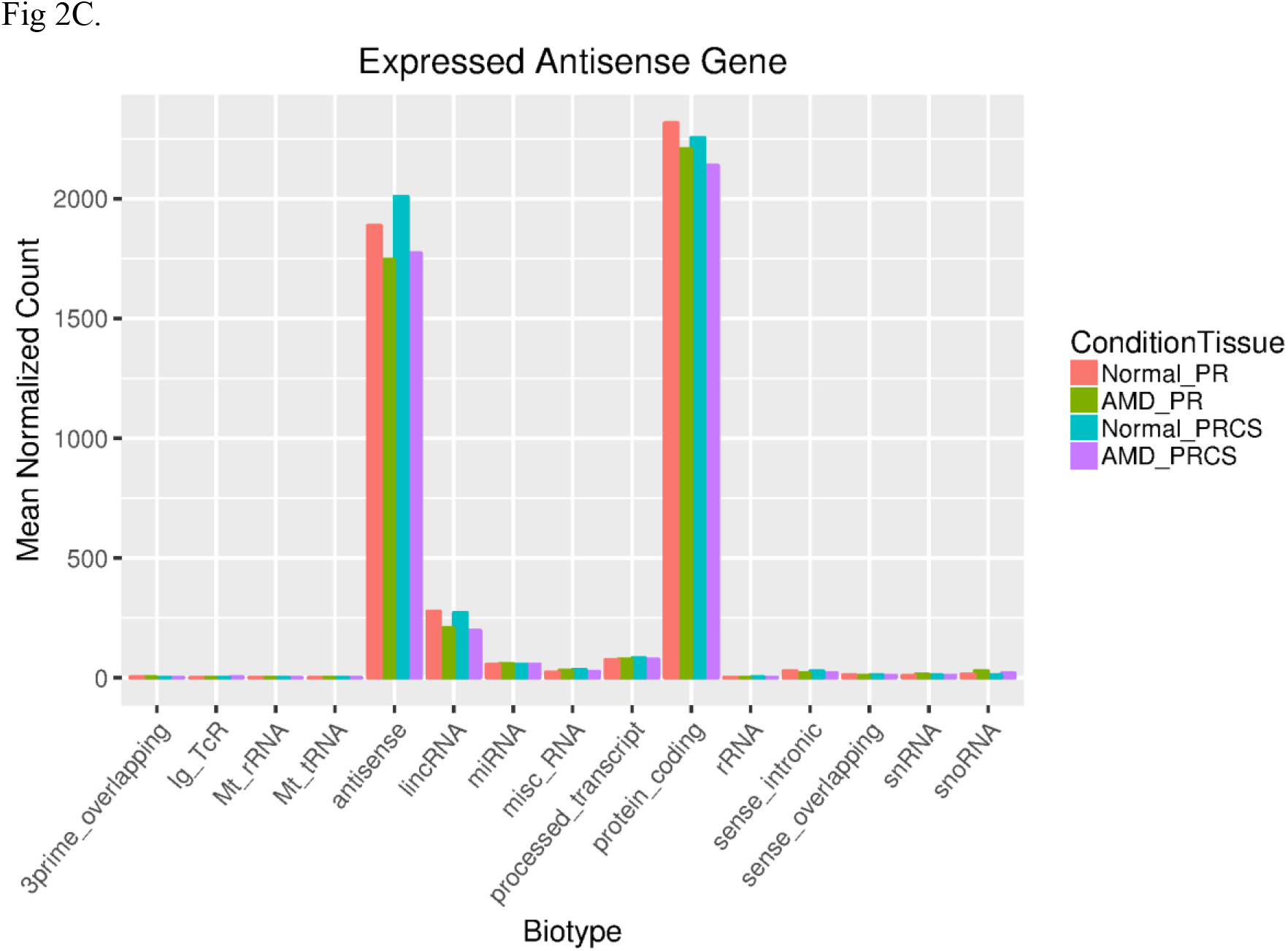
Distribution of biotypes of expressed transcripts in each category of biotypes is plotted by condition and tissue type. A) All expressed sense transcripts, B) Expressed ncRNA and c) Expressed antisense transcripts.

To explore the differences between the retina and PRCS tissue types and between normal and AMD eye tissues, we performed hierarchical clustering of the top 1,000 most variably expressed ncRNAs. These were defined as those with the largest coefficient of variation of PORT normalized average coverage (Fig.3). As expected, we observed a clear delineation between the PR and PRCS samples. Following an emerging trend, the delineation of expression between normal and AMD PR, as well as normal and AMD PRCS samples was apparent. Our results indicate differential expression profiles or specialized signatures for transcripts between PR and PRCS in the normal and diseased states.

**Fig. 3.**
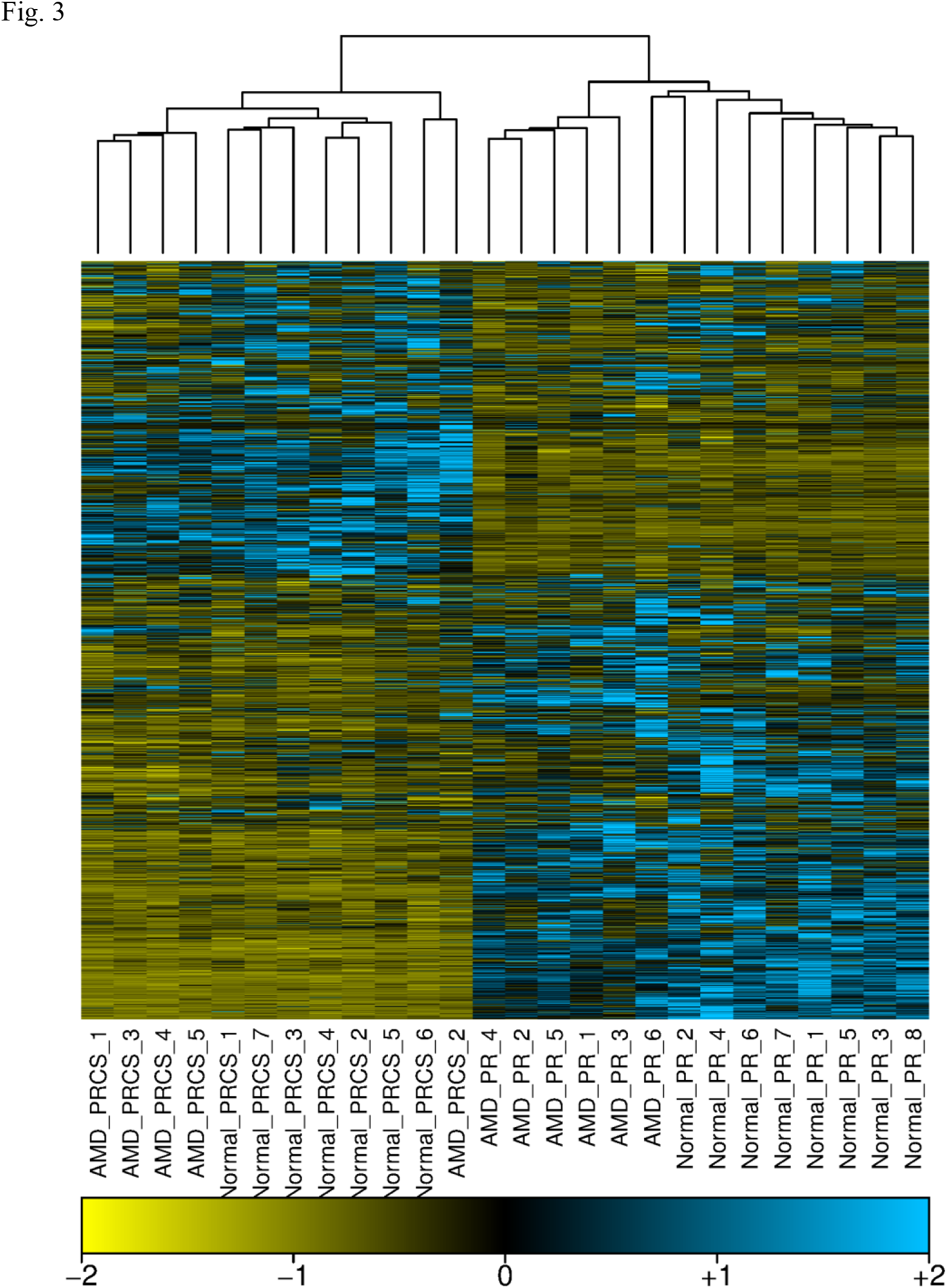
Expressed ncRNAs with wide variability. Heatmap of expression in average coverage of the top 1000 most variably expressed ncRNAs, defined as those with the largest coefficient of variation across all tissue/disease types.

### Differential anti-sense gene expression in AMD

An extreme differential expression of anti-sense genes transcripts was observed in the AMD-PR and AMD PRCS tissues when compared to the normal tissues. There are 2,025 anti-sense transcripts up-regulated and 597 transcripts down-regulated when normal PR was compared to the AMD PR (q=0.0001). Comparison between normal and AMD PRCS resulted in 941 upregulated and 510 down regulated antisense genes that were differentially regulated (q=0.001). Figure 4 shows a typical example; the single-exon gene RN7SK on Chromosome 6 is highly expressed in retina, as shown in the red depth-of-coverage plot. While there is a trace amount of anti-sense transcription of RN7SK in normal controls, as shown in the blue depth-of-coverage plot at the bottom, there is an high anti-sense transcription of this gene in AMD, as shown in the blue depth-of-coverage plot at the top. This is a typical example among thousands of genes, which strongly targets a number specific pathways and networks. We observed this in the PRCS tissue as well.

Although there are hundreds of genes with down-regulated anti-sense transcription, they do not tend to target particular pathways. In contrast, among those genes with upregulated anti-sense transcription in the retina, the EIF2 signaling pathway was predominant (enrichment *p*-value < 3.31E-26), suggesting a central role for this pathway implicating ribosomal upregulation. Other pathways that were significantly affected were regulation of elF4 and p70S6K signaling, mTOR signaling, phototransduction and mitochondrial dysfunction pathway. The antisense transcripts that were upregulated also appear to manage apoptosis pathways, mitochodrial function and NRF2 mediated oxidative stress response, which is known to contribute to retinal maintenance.

The top networks identified in our analysis include cell death and survival, cellular growth and proliferation and cellular assembly and organization as well as RNA posttranslational modification, repair and connective tissue disorders. The top relevant toxicology lists were mitochondrial dysfunction, NRF2-mediated oxidative stress response and PPAR/RXR RXR activation pathways. In addition to the pathways affected in the retina are the EIF2/mTOR, the differentially expressed antisense RNA in PRCS tissues on IPA analysis included those involved in clathrin and caveolar mediated endocytosis signaling, actin cytoskeleton signaling and RhoGDI signaling. Cellular function and maintenance, cellular development, cell death and survival were key molecular and cellular functions along with nervous system and development function, which are regulated in PRPC.

Among individual upregulated anti-sense transcripts in the retina, we found nearly five-fold higher expression of Transferrin in the AMD retinal tissue when compared to the normal retina (*q*-value=9.68 E-16). Many other protein coding genes up regulated in AMD include glyceraldehyde-3-phosphate dehydrogenase, nuclear ubiquitous casein and cyclin-dependent kinase substrate 1(NUCKS1), glutathione S-transferase alpha 4 (GSTA4) and interphotoreceptor matrix proteoglycan-1 (IMPG1), a gene encoding the Sialoprotein associated with cones and rods (SPACR) ^30^. The expression of many small nuclear pseudogenes and small nucleolar RNA such as SNORD3A and SNORA73B appear to be upregulated in AMD, implying a global regulation of transcription machinery necessary for the normal retinal maintenance is affected in AMD (Supplementary Table 2). The lincRNA RMRP (RNA component of mitochondrial RNA processing endoribonuclease) is the predominant lincRNA that is significantly overexpressed in the AMD retina when compared to the normal tissue. Although it is known to be expressed in mouse and human tissues and implicated in early murine development, its role in the retina has not been established ^31, 32^. Among the DE antisense transcripts, we identified matrix remodeling associated 8 (MXRA8), prickle planar cell polarity protein 4 (PRICKLE4) and many solute carrier family proteins among other protein coding genes which appeared to be significantly downregulated at least 3 to 4-fold in AMD retina when compared to the normal retinal tissue.

We identified several antisense transcripts in the PRCS tissues that are significantly differentially expressed in AMD PRCS when compared to the normal PRCS tissues. The antisense transcription of protein coding genes EEF1A1, COL8A1, EEF2, APOD, RPE65, CLU, UBC and other ribosomal proteins responsible for eukaryotic transcription was selectively upregulated 3 to 4-fold in the AMD PRCS when compared to the normal PRCS (q values > 1 E-15). Many noncoding transcripts mostly belonging to the snoRNA family such as SNORD3A, SNORA73B and SNORD17 were upregulated 3 to 4-fold more in AMD PRCS tissues when compared to normal PRCS tissue. The predominant antisense transcripts that appeared significantly downregulated (over 3 fold, q value >1E-12) in AMD PRCS when compared to normal PRCS include RBP5, MST1, MYL5 and LCAT among other lincRNA, indicating that antisense transcription is highly varied in both retina and RCS tissues during age-related macular degeneration.

### Differential sense expression in AMD

For the Normal PRCS vs AMD PRCS comparison, the *q*-value cutoff of 0.25 was used for both the sense genes and for the ncRNA, identifying 310 and 188 differentially expressed genes (DEGs) respectively. A total of 537 and 280 DE transcripts (q-value cutoff of 0.05) were identified in the Normal PR vs AMD PR comparisons for sense genes and ncRNA tissues respectively. Table 1 details the results and Fig. 5A displays the expression of those DEGs across conditions. The difference between tissues was larger than the difference between disease states within the same tissue indicating a tissue specific expression profile of ncRNA. It is also interesting to note that there are transcripts differentially expressed (DE) between normal and AMD regardless of tissue. It is unclear whether this may be a result or cause of AMD progression. As we are interested in the effect of disease on the noncoding landscape, we also determined the biotypes of all DE ncRNAs (Fig. 5B).

**Fig. 4.**
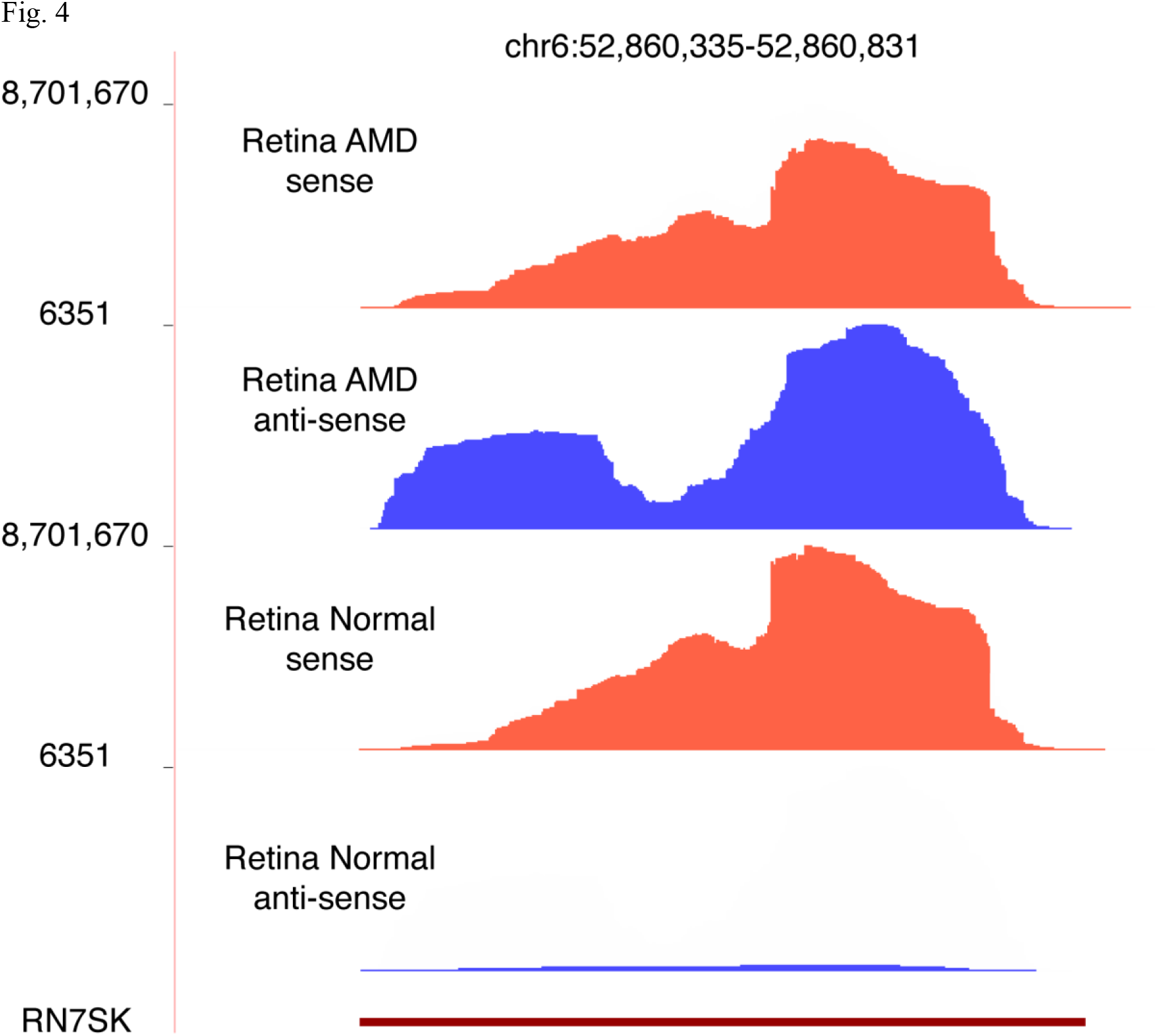
Extreme differentially expressed anti-sense transcription (in blue) of the RN7SK gene. The coverage plot was generated merging normalized coverage of 5 replicates in each condition. Sense gene counts of five AMD samples were: 2,245,429, 2,029,045, 2,326,472, 1,986,731, 1,836,122; Sense gene counts of five Normal samples were: Antisense gene counts of five AMD samples were: 2366, 2359, 2084, 1128, 1583; Antisense gene counts of five Normal samples were: 51, 41, 82, 44, 102.

**Fig. 5.**
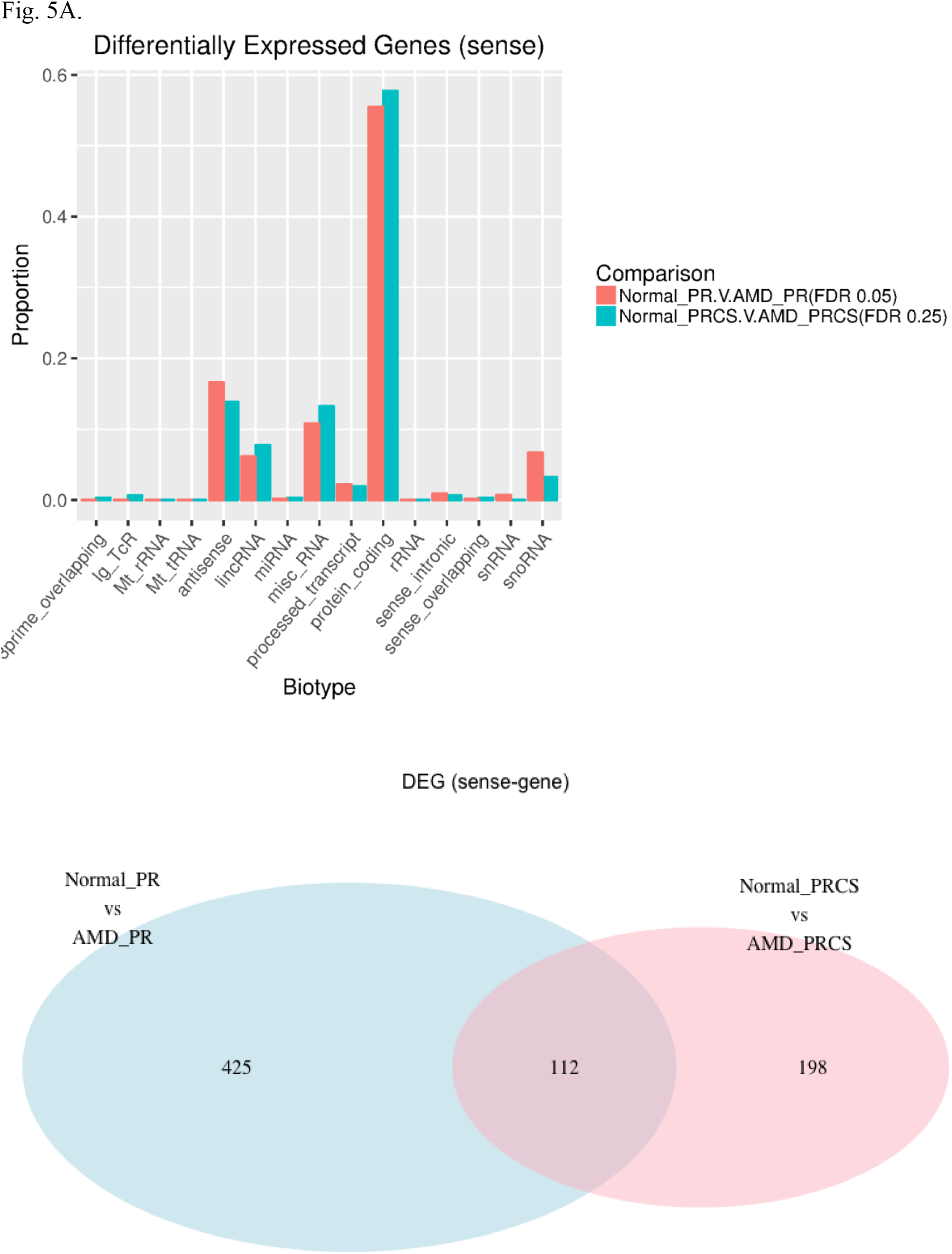
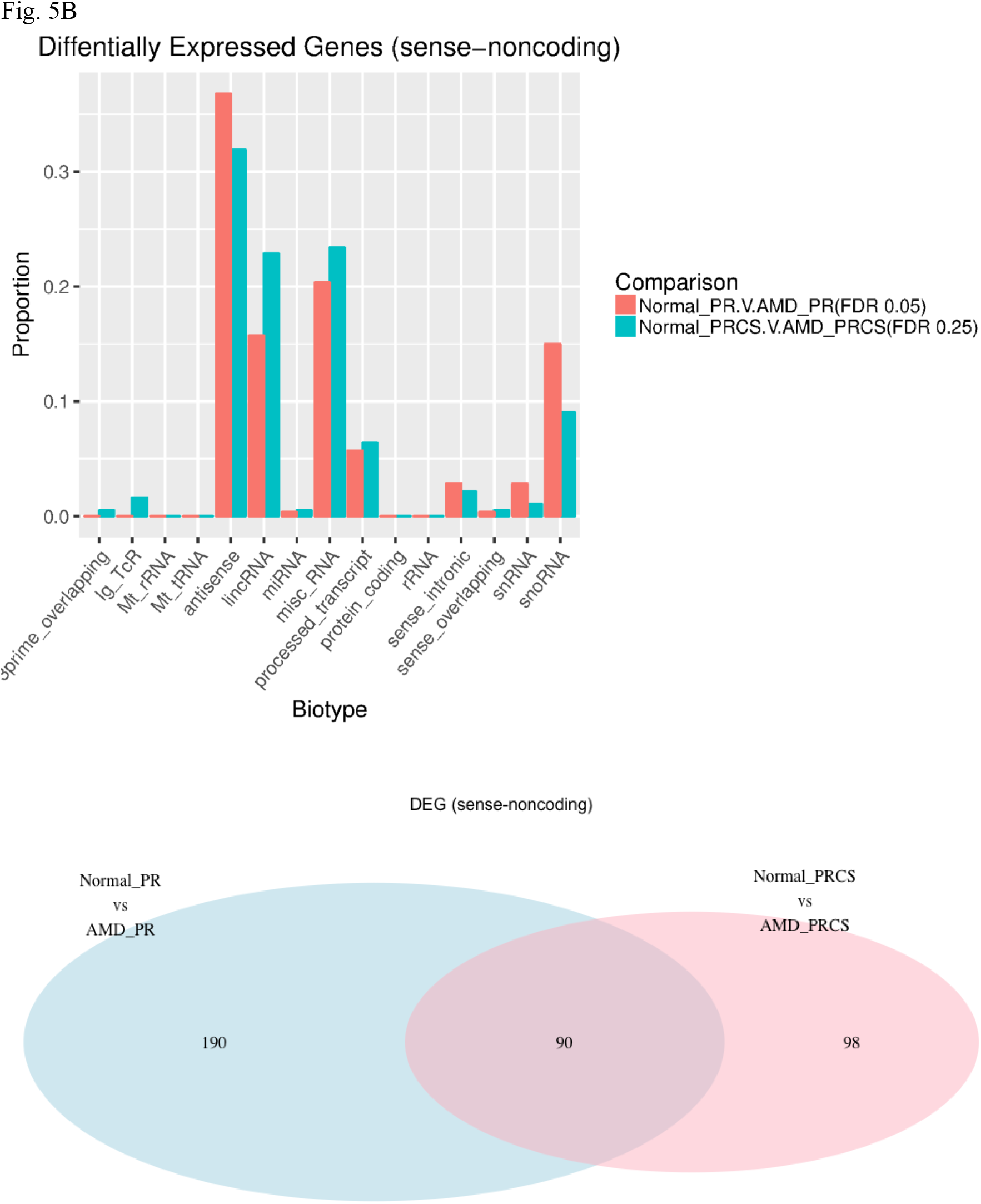
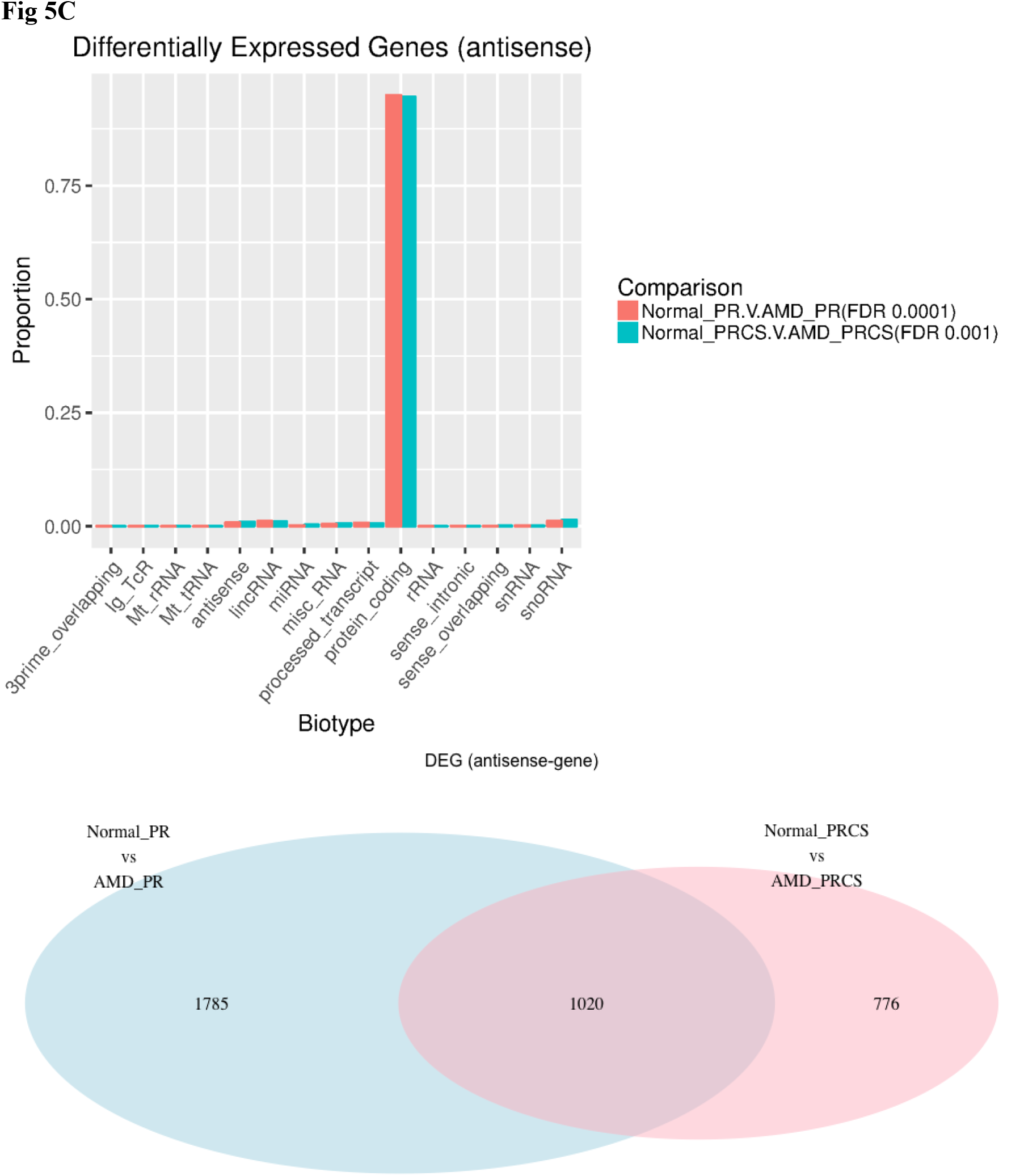
Differentially expressed transcripts between normal and AMD comparisons were represented as bar graphs and Venn diagrams between. (A) PR and PRCS sense transcripts (B) PR and PRCS noncoding RNA and (C) PR and PRCS antisense RNA transcripts.

**Table 1.** The number of differentially expressed noncoding genes

| Comparison of disease states in tissues | Biotype | Number of DE transcripts |
| --- | --- | --- |
| Normal PR vs AMD PR | Gene antisense | 2622 (2025 up; 597 down) |
| Normal PRCS vs AMD PRCS | Gene antisense | 1451 (941 up; 510 down) |
| Normal PR vs AMD PR | Gene sense | 537 (211 up; 326 down) |
| Normal PRCS vs AMD PRCS | Gene sense | 310 (176 up; 134 down) |
| Normal PR vs AMD PR | ncRNA | 280 (170 up; 110 down) |
| Normal PRCS vs AMD PRCS | ncRNA | 188 (136 up; 52 down) |

### Co-localization of DE-ncRNA and pathway analysis of nearby protein-coding genes

Since most ncRNA are not well characterized and some may be involved in regulation of their neighboring coding-genes, we looked at protein coding *genes in cis* to our DE ncRNAs. To find these neighboring genes, we used GREAT with default settings and imported these co-localized coding genes into Ingenious Pathway Analysis (IPA) (Qiagen) for further analysis. All AMD GWAS associations were found using the NHGRI-EBI Catalog and lifted over from GRCh38 to hg19 using Ensemble’s Converter tool. The DE genes, ncRNA and antisense genes were separately investigated for their role in the different pathways that may affect the retina and RCS function.

The majority of the noncoding RNA that we found to be differentially expressed in the AMD PR vs Normal PR and AMD PRCS vs Normal PRCS belonged to antisense transcripts followed by small nucleolar RNA and a few lincRNA that were significantly upregulated by over 3 to 4 fold. Most of these noncoding RNA were either novel or have uncharacterized function; their role in the eye is yet to be explored. The DE ncRNA from both the retinal and PRCS tissues were co-localized with the entire gene network using GREAT software analysis and the resulting genes were analyzed using IPA pathway analysis tool to understand pathways that are regulated by DE ncRNA. IPA analysis of GREAT genes from the retinal and PRCS tissues identified Epithelial Adherens Junction and corticotrophin releasing harmone-signaling pathways as the predominant pathways being regulated by the ncRNA. While the retina specific analysis identified Gap junction signaling and germ cell signaling pathways as top canonical pathways, the RCS tissue analysis revealed remodeling of epithelial adherens junction, dermatan sulphate biosynthesis (late stages) and axonal guidance signaling as major pathways regulated by the ncRNA.

IPA analysis of the sense gene transcripts differentially expressed in the AMD vs normal retina tissue revealed IL-22 signaling, amyloid processing, 4-1BB signaling in T lymphocytes as the top canonical pathways. Similar analysis in PRCS tissues revealed IL-22 and IL-10 signaling as the top canonical pathways along with Cardiolipin biosynthesis II and role of JAK family kinases in IL-6 type cytokines signaling pathways. The immune related pathways that were obtained in the IPA analysis in both tissues are very relevant in causing AMD pathobiology as increased circulation of complement component 5a (C5a) was observed in serum circulation of AMD patients which increases the expression of IL-17 and IL-22 cytokines ^33^. It was also reported that anti-inflammatory cytokine IL-10 have both an increased accumulation of macrophages in neovascular lesions and decreased choroidal neovascularization thus causing retinal pathology ^34^.

### Discussion

In this study we present the first transcriptome data analysis that compares retinal and RCS tissues isolated from human cadaver normal and AMD eye globes. While differential expression between gene transcripts and ncRNA have been observed between the retina and RCS tissues in the normal and AMD tissues, we have also observed intriguing differences between the control and AMD tissues. For the first time, we report an extreme increase in anti-sense transcription in AMD eye tissues, implicating a potential role in pathology of AMD. Our study mainly focused on understanding the differential expression of selected genes, non-coding and antisense transcription in PR and PRCS tissues that emerged from our RNA-Seq analysis.

Antisense transcription is known to regulate the gene expression either *in cis* or *in trans,* either through transcription or by association with non-coding RNA in the genome ^35^. The presence of antisense transcripts can induce a threshold-dependent regulatory control to fine-tune gene expression. It is previously reported that budding yeast increases its gene expression between different cells by managing the balance between sense-antisense transcripts. ^37^ The stress related genes were enriched for antisense transcripts in yeast. ^37^ Using our bioinformatics pipeline, we successfully analyzed and established the RNA transcripts produced for sense and anti-sense transcription encoded by the same genomic region. This may be useful to characterize both sense and anti-sense regulatory mechanisms employed by tissues.

Hierarchical clustering of our samples and diseased tissues was appropriately performed and clean separation was achieved only when we used anti-sense genes for clustering - this is unique and not attempted to date in RNA-Sequencing studies. The differential expression of anti-sense transcripts between normal and AMD retina and normal and AMD RCS tissues revealed eIF2 regulation as the predominantly upregulated pathway in AMD tissues. The phosphorylation of eIF2alpha at Ser51 is known to inhibit the translation initiation causing a temporary shutdown of protein synthesis^38, 39^. Persistent eIF2 alpha phosphorylation through regulatory kinases is reported during stress conditions in neurodegenerative diseases like Alzheimer’s disease (AD) ^40^. The phosphorylation of eIF2 is controlled by four kinases among which PKR-like endoplasmic reticulum kinase (PERK) along with GCN2, HRI and PKR^40^. Interestingly, the PERK/eIF2α/ATF4 and IRE1/ASK1/JNK cascades are the most important pathways that were associated with pathological changes like inflammation, ER and mitochondrial stress and matrix degradation in AMD ^41, 42^. These pathways can elicit several AMD-related pathological changes via the induction of VEGF, C/EBP homologous protein (CHOP), caspase-4 (CASP4), and nuclear factor-κB (NF-κB) ^43^ The differential up regulation of eIF2 observed in AMD retina and RPE may be due to increased stress response and inflammation.

Differential antisense expression of key complement genes in the RCS includes C1R, C3, CFH, which are upregulated more than two fold in the AMD tissues when compared to the normal tissues. The antisense RNA for other key apolipoproteins known to play a key role in AMD, such as APOE and APOE, were also differentially expressed in AMD tissues. These genes are critical for maintaining RPE and retinal homeostasis, and disruption of these processes may lead to AMD. The magnitude of differential anti-sense expression observed in our study suggests that anti-sense transcription could provide another level of gene regulation in addition to post-translational and transcription factor-mediated mechanisms.

When comparing the total DE ncRNA profiles, we observed a predominant expression of small nucleolar RNA (snoRNA) in AMD tissues when compared to normal tissues. Among the DE snoRNA, we found 3-fold higher expression of SNORA73B, SNORA54 in both retina and RCS tissues of AMD when compared to the normal tissues, indicating that they may have a role in AMD. The DE ncRNA also identified eIF2-signaling pathways as the key pathways in IPA analysis for both the retina and PRC tissues reiterating their role in AMD.

We employed strict thresholds for minimizing false positives when interpreting the differentially expressed genes, non-coding and gene antisense RNA and compared expression between PR and PRCS tissues for each sample in both control and AMD tissues to validate our results. Second, due to the limited RNA availability of the macular retina and RCS tissues in the AMD donor eye samples (a region which is necessary for central vision and critical in age-related macular degeneration), our study only addressed the differential expression and transcriptome changes observed in the peripheral retina and RCS tissues. The sample size (n=8 donors) gave us enough power to detect transcriptome level differences between the normal and AMD tissues analyzed in our study. Nevertheless, transcriptome analysis of a large dataset of samples with different AMD progression levels and types (early to advanced/ neovascular forms) is warranted to identify stage specific expression of DE non-coding RNA and genes that may be used as biomarkers for tracking AMD progression and pathology. Our data also showed strong correlation with the previously published FPKM values of the peripheral retina and RCS tissues reported previously^44,45^.

Pseudogenes dominated the differentially expressed transcripts in both the Retina and RCS tissues in our study. We found many common pseudogenes that were differentially upregulated and down regulated in both of these tissues. As their roles in biology and human disease and eyes are not fully investigated, we have removed them from our analysis to focus on the remaining signatures, potentially eliminating some pseudogenes that may be functionally important during AMD.

With the advancement of next generation sequencing technologies, increased depth in sequencing transcriptome and reduced costs, there is a promise that more eye-specific data will be available to analyze the differential non-coding transcriptome profiles in individual eye tissue layers. This will provide an opportunity to uncover novel pathways to study the pathophysiology of different eye diseases such as retinal degeneration, macular degeneration and other eye pathologies regulated by ncRNA. In summary, the present transcriptome analysis in the retina and RCS tissues greatly increased our knowledge of the entire coding and non-coding regions of the genome expressed in these tissues. However, the exact spatial expression patterns of most of these genes and ncRNA are still unclear, as are the *in vivo* functions of these ncRNAs in retinal/ocular development and AMD pathogenesis. Functional studies of ncRNAs in the retina and other ocular tissues have the potential to greatly enrich our understanding of normal and disease processes of the eye and inspire novel therapeutic strategies.

## Materials and Methods

### Eye Collection

Our study conformed to Institutional Review Board regulations for use of human tissues at University of Alabama and at University of Pennsylvania (Penn). We have utilized eye tissues collected from two sets of eight pairs of eyes from non-diabetic Caucasian donors with a death-to-preservation interval of < 6 hr. The first set is collected from donors with mean age of 73.9 yr ± 12.5 yr (mean ± standard deviation), and, the second set was collected from donors with mean age of 84.6± 7.2 yr (mean ± standard deviation), to maximize the number of eyes with AMD pathology. All eyes were collected and processed by the Alabama Eye Bank recovery personnel and are preserved in RNA-later (Qiagen, Valencia, CA, USA) for the left eye and, 2% glutaraldehyde and 1% paraformaldehyde in 0.1M phosphate buffer for the right eye ^21^. The left eyes were shipped overnight on wet ice to Penn, and were processed upon arrival. All the eye tissues for our study have been obtained from tissues collected by Dr. Dwight Stambolian and Dr. Christine A. Curcio as a part of the Arnold and Mabel Beckman Initiative for Macular Research foundation grant (2014). The normal or AMD status of donor eyes were assessed by Dr. Christine A. Curcio at the University of Alabama by a three-component protocol as described previously ^21^. The fellow eye design was adapted in our study following well-documented literature ^46-48^. The left eyes preserved in RNA-later solution were examined by photography with stereomicroscope before dissections. For each donor eye, the retina (PR) and RPE-Choroid-Sclera (PRCS) samples were dissected from the peripheral region of the posterior eye globes using a 10mm-biopsy punch followed by a 8mm-punch in the middle of the 10mm-punch to minimize the sample contamination. The PR was collected separately from PRCS into a 1.5ml tube and stored separately until further processing.

### Library preparation and RNA-Seq runs

From 8 normal and 8 AMD donor eyes are a total of 32 RNA samples were prepared from 16 PR and 16 PRCS samples using the AllPrep DNA/RNA Mini kit (Qiagen). The RNA quality was determined using R6K Screen Tape on a 2200 Tape Station (Agilent, Santa Clara, CA, USA) and was quantified using Qbit-BR (Broad Range) assay kit on a Qbit 2.0 Fluorometer (Life Technologies) following manufacturers instruction. Only RNA with integrity number (RIN) value of >8.5 was used for preparing sequencing libraries. To sequence the transcriptome, library preparation and sequencing was done using the TruSeq Stranded total RNA with RiboZero Gold kit (Illumina, CA) protocol with a total RNA (800ng) as the starting material. A total of 32 libraries were prepared with unique barcode sets and their quality was determined using Agilent DNA1000 chip following manufactures protocol. All the DNA libraries with mean peak size of 260bp were processed for sequencing. The libraries were sequenced on an Illumia HiSeq 2000 machine following manufacturers protocols. A total of 16 lanes were run for sequencing (2 libraries/lane with a 100bp Paired End reads) all 32 libraries and to achieve sequencing depth of 200 million reads per sample (Supplementary Table 1).

### RNA-Seq quality control

The RNA-seq data for all the samples in our study are deposited and can be accessed with Geo accession number GSE99248. Each sample is sequenced to a depth of around 200 million 100-bp paired-end reads per sample (Supplementary Table 1). Pre-alignment QC showed average quality scores for both forward and reverse reads to be >=30 throughout the length of transcripts. Reads were then mapped to hg19 using STAR ^49^ with a mapping rate >=93% for all samples. Post-alignment QC revealed one Normal RPE/Choroid sample with an extremely high level of rRNA, two AMD RPE samples with detectable levels of retinal contamination, and three AMD samples (two retina and one RPE samples) with high chromosome M percentage. These six samples were removed prior to further analysis (Supplementary Table 1). Two samples (TR01 and TR13) were resequenced owing to decreased number of reads obtained for each sample by running them individually on each lane on HiSeq2000.

### RNA-Seq Data analysis

RNA-Seq reads from each sample were aligned to hg19 using STAR version 2.5.1b ^49^. Data were normalized at the read level, prior to quantification, using a resampling strategy PORT v0.8.2a-beta (https://github.com/itmat/Normalization). All pseudogenes were filtered from the ENSEMBL annotation (GRCh37.p13) due to unreliable alignments. The normalized SAM files were then quantified at the gene level by identifying, for each gene, all reads that were consistent with some ENSEMBL annotation splice form of the gene. Differential expression (DE) analysis was performed between each pair of tissue/disease type by calculating Limma-Voom p-values for each gene and then performing a Benjamini-Hochberg correction for multiple testing, to produce q-values ^50^.

### Gene Expression analysis

The overall gene expression (including coding and noncoding transcripts) was obtained for each sample using PORT. Hierarchical clustering based on Jensen-Shannon divergence was performed in R ^51^.

### Noncoding RNA identification, classification, and quantitation

Transcript biotypes were defined using the “gene biotype” information in the ENSEMBL annotation (GRCh37.p13). Those ncRNA with average depth of coverage equal to, or greater than 1 were considered expressed.

### Co-localization analysis

The genomic coordinates of DE ncRNAs were imported to the Genomic Regions Enrichment of Annotations Tool (GREAT) with default parameters for co-localization analysis ^52^. For pathway analysis, Ingenuity Pathway Analysis (IPA) software (Qiagen, CA) was used.

## Acknowledgements

The authors thank Stephanie Yee for her help with the eye dissections. The authors also thank the financial support received from BrightFocus Foundation grant (VRMC), Research to Prevent Blindness Unrestricted Grant Funds to Scheie Eye Institute (VRMC), F.M. Kirby Foundation, and The Paul and Evanina Bell Mackall Foundation Trust, The National Center for Advancing Translational Sciences (5UL1TR000003, Garret A. FitzGerald).

